# The Role of Environmental Information and the Limitations of Predictability with Transformer-based Time-Series Models for complex Astrocyte Trajectories

**DOI:** 10.64898/2026.10.15.759300

**Authors:** Koya Takahashi, Kaito Zama, Yohei Shibata, Sora Endo, Hidenori Tabata, Kazunori Nakajima, Noriko F. Hiroi

## Abstract

Morphogenesis is the process by which an organism develops from a single cell into a complex organism, and one factor contributing to this complexity is the interaction between cells and their environment. In heterogeneous environments comprising differing cell types and polymeric structures, the observed complexity appears to approach an infinite value. Recent developments in imaging tools and deep learning for image analysis enable the extraction of meaning from terabyte-scale images at a realistic computational cost. In this paper, we used a Transformer-based model to predict the migration trajectories of astrocytes during mouse brain development, aiming to quantitatively elucidate (i) what makes prediction difficult, i.e., the structure of predictability and its theoretical limit, and (ii) to what extent environmental information (vascular structure) and multimodal modeling of the predictive distribution can improve prediction. Based on a bivariate Gaussian distribution (*µ, σ, ρ*) at each time point, we predicted future positions while accounting for uncertainty and evaluate the quality of the predictive distribution. We report three main findings. First, point-prediction accuracy reached a fundamental limit set by the irreducibility of the movement direction; the error curve was described with high accuracy by the mean-squared-displacement law of a persistent random walk. Second, extending the predictive distribution from a single bivariate Gaussian to a continuous mixture (mixture density network, MDN) captured the run/rest bimodality without discretization and substantially improved the quality of the predictive distribution (negative log-likelihood, NLL) while preserving point-prediction accuracy. Third, a model that integrated vascular environmental information into this multimodal model achieved the best performance in point-prediction accuracy, consistently across three training seeds, showing that environmental information contributes to prediction in combination with multimodal output. This study presents both an improvement of the predictive distribution for complex cell migration and a framework for quantifying the limit of predictability based on movement statistics.

## 1. Introduction

Transformer is a deep learning architecture originally proposed by Vaswani et al. [1] with the aim of improving performance in machine translation tasks within natural language processing. Its most distinctive feature is that it completely eliminates the recursive structure found in traditional RNNs (Recurrent Neural Networks) and instead adopts a self-attention mechanism that simultaneously calculates the relationships among all elements in the input data. This structure makes it possible to capture long-range dependencies within a sequence extremely effectively. Furthermore, because it enables a high degree of computational parallelisation, it is considered excellent for efficient learning using large-scale data. On the other hand, several challenges have been identified regarding the processing of time-series data. In particular, the computational complexity and memory usage of the self-attention mechanism increase quadratically (*O*(*N*^2^)) with respect to sequence length *N*, making it difficult to process very long sequences [2]. Additionally, because the architecture relies solely on positional encoding, it is unable to fully capture local and global temporal structures specific to time series—such as continuous temporal changes, periodicity, and trends—which are cited as its main areas of challenge [3].

Astrocyte migration during brain development exhibits extremely complex behaviour, and predicting their future trajectories and positions is one of the most challenging problems in biomedicine. To comprehensively elucidate tissue morphogenesis processes, it is necessary to capture the temporal changes in the orchestration of the movement of cells; however, the sheer volume of time-lapse data and the irregularity of cell behaviour have posed serious computational bottlenecks for conventional analytical methods. In response to this challenge, the recent emergence of advanced deep learning systems has opened up new avenues of possibility for predicting complex cell trajectories over time using data-driven approaches.

However, applying a standard time-series Transformer model directly to astrocyte trajectory prediction presents challenges. The complex movement and rapid morphological changes of cells involve fluctuations in local temporal structures, which conventional models struggle to capture. Furthermore, cell migration behaviour is not determined in isolation; instead, it is supposed to depend on information about the surrounding environment—such as vascular structures—and local spatial features contained in cell-tracking images. To capture these intricate spatiotemporal dynamics, we use a hybrid model integrating a Convolutional Neural Network (CNN) and a Transformer. While the CNN excels at effectively extracting local spatial features from image data (e.g., vascular structure and microenvironmental contexts) [4,5], the subsequent Transformer captures the long-range temporal dependencies. By feeding the CNN-extracted spatial features into the Transformer through time-aligned cross-attention, this method makes predictions that inherently account for spatial context. In addition, we note that cell migration has a multimodal nature, intermittently switching between movement (run) and rest, and we therefore extend the predictive distribution from a single Gaussian to a continuous mixture of Gaussians (mixture density network, MDN) [6]. Multimodality of prediction was originally addressed for AIS ship trajectories by TrAISFormer [7] through spatial discretization; however, because such discretization breaks down for the small displacements at the cellular scale [8], we instead recover multimodality in a continuous displacement space.

In this study, we applied this spatiotemporal framework to time-lapse data of astrocytes during actual brain development and obtained the following three key findings. First, we showed that point-prediction accuracy reaches a fundamental limit set by the irreducibility of the movement direction. This limit is described with high accuracy by the mean-squared-displacement (MSD) law of a persistent random walk, and we further quantified—information-theoretically—that the predictability of astrocyte migration is far lower than that of human mobility [9]. Second, we showed that extending the predictive distribution from a single bivariate Gaussian to a continuous mixture (MDN) captures the run/rest bimodality without discretization and substantially improves the quality of the predictive distribution (negative log-likelihood, NLL) while preserving point-prediction accuracy; almost all of this improvement arises in the transition from a unimodal to a bimodal distribution, revealing that the predictive distribution is essentially bimodal. Third, a model that integrates vascular environmental information into this multimodal model achieved the best performance in point-prediction accuracy, consistently across three training seeds, showing that environmental information contributes to prediction in combination with multimodal output. This study not only improves the predictive distribution for complex cell migration but also presents a new theoretical framework for quantifying the limits of predictability based on movement statistics.

## 2. Materials and Methods

### 2.1. Time-lapse imaging of mouse brain slices

ICR mice (Japan SLC, Shizuoka, Japan) and Flt1-DsRed mice [10] were housed at Institute for Developmental Research. All mice were kept at 22-24°C with 40-60% humidity under a 12-hour light/12-hour dark cycle. At noon on the day a vaginal plug was observed was considered embryonic day 0.5 (E0.5). Male heterozygous Flt1-DsRed mice were bred with female wild-type (ICR) mice (both 10-24 weeks old). At embryonic E15, *in utero* electroporation was performed as described previously[11–13] to introduce the rDIO system [14] (pCAG-hyPBase [15], pPB-CAG-roxed-DIO-EGFP, hGFAP-Cre [16], pDcx-Dre [17], 0.5 *µ*g/mL each) into the ventricular zone of the embryonic cerebral hemispheres.

Two days later (E17), time-lapse imaging of the mouse brain slices was performed as reported in the previous studies [14,18,19]. Briefly, the brains were dissected and embedded in 3% low melting point agarose (SeePlaque; Lonza, ME, USA) in Hanks’ balanced salt solution (HBSS; Wako, Japan) containing 0.14 g/L CaCl_2_ and 0.2 g/L MgSO_4_·7H2O. Next, the brains were sectioned coronally at 250 *µ*m using a vibrating microtome (VT1000, Leica, Germany). Brain slices obtained from around the level of the interventricular foramen were placed on a Millicell-CM (pore size, 0.4 *µ*m; Millipore, Germany) in a glass-bottomed dish (Iwaki, Japan) containing Neurobasal medium (Thermo Fisher Scientific, MA, USA) with 10% fetal bovine serum (FCS), 2% B27 (Life Technologies), 50 units/mL penicillin, and 50 *µ*g/mL streptomycin(Thermo Fisher, USA). The culture dish was then placed in an incubator chamber (5% CO_2_ and 40% O_2_ at 37°C, Tokai Hit, Japan) fitted onto a confocal microscope (A1R; Nikon, Japan).

Approximately 8 to 15 optical Z sections for 6 to 12 positions were obtained automatically every 15 min, and about 10 focal planes (*∼* 50 *µ*m thick, 20*×*, 0.50 N.A., Nikon) were merged to visualize the entire blood vessel and cell trajectories. The movie files were constructed using ImageJ software (National Institutes of Health, [20]). Figure 1 shows examples of 8 random image data sets.

**Figure 1.**
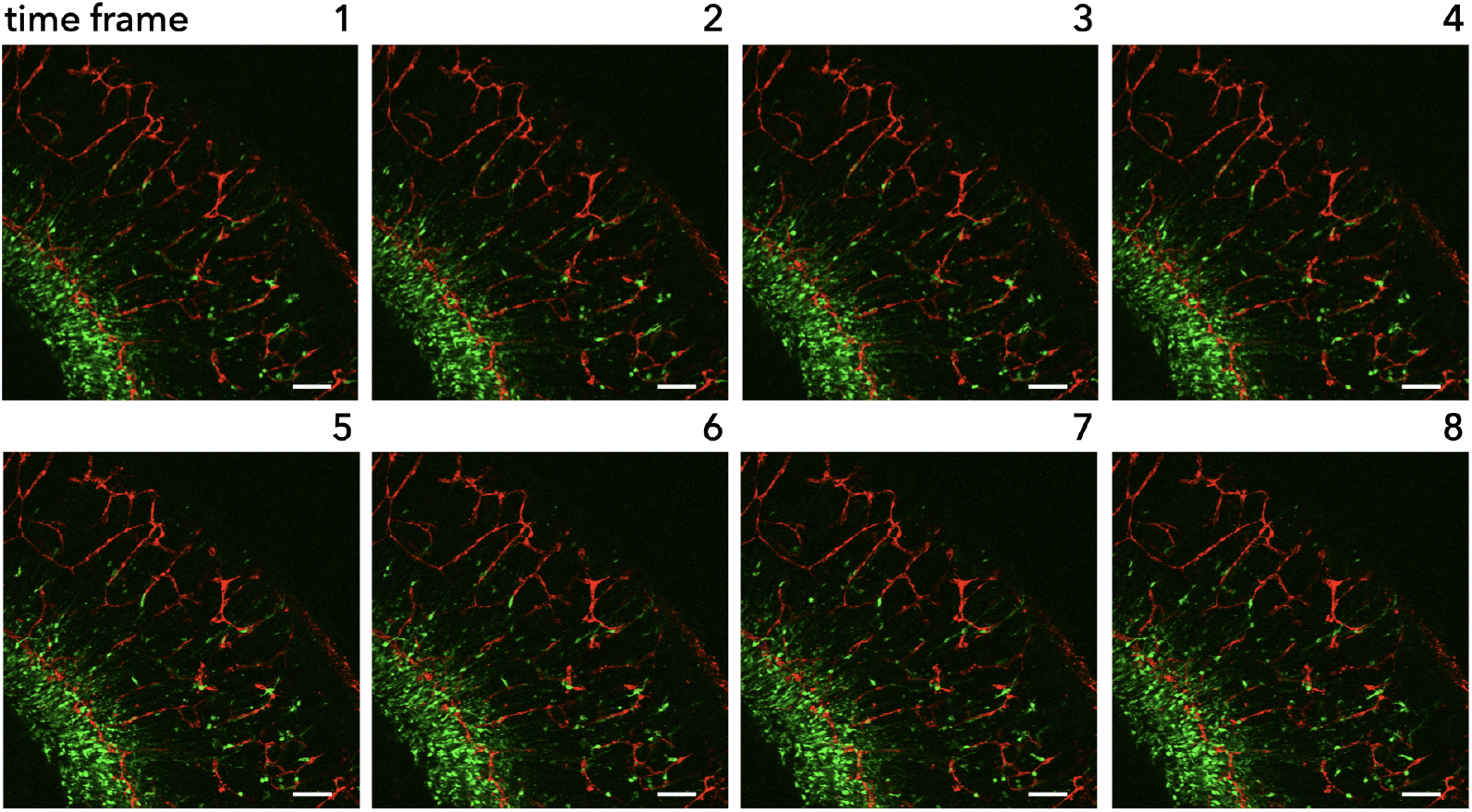
A selection of sliced brain images. Images were taken every 15 min. Astrocytes: green, blood vessels: red. Bar = 100 *µ*m

### 2.2. Image data processing

Cell coordinates, velocity, and direction were extracted from the movie data using Fiji [21] together with the TrackMate plugin [22,23]. Tracking with a DoG (Difference of Gaussian) detector generated a total of 30,521 trajectories.

The distribution of track lengths is shown in Figure 2. The distribution is strongly skewed toward short tracks (median 18 frames), containing many short trajectories. To ensure a length sufficient for analysis, only trajectories of 20 time points (frames) or longer were retained. This threshold left 14,417 trajectories, which were split into training, validation, and test sets in a 70%/20%/10% ratio (Table 1). For each trajectory, the first 10 time points were given to the model as context, and positions from the 11th time point onward (up to a maximum sequence length of 150) were predicted. Velocity and direction were computed from positional differences. Each feature was min–max normalized within a prescribed range, and variable-length trajectories were padded up to the maximum sequence length with masking.

**Table 1.** Dataset statistics and split. Of all tracks obtained by tracking, those meeting the minimum-length threshold were split 70/20/10.

| Item | Value |
| --- | --- |
| Total tracks obtained by tracking | 30,521 |
| After minimum-length threshold ($\geq 20$ frames) | 14,417 |
| Training (70%) | 10,091 |
| Validation (20%) | 2,883 |
| Test (10%) | 1,443 |
| Context length (init_seqlen) | 10 time points |
| Maximum sequence length (max_seqlen) | 150 time points |

**Figure 2.**
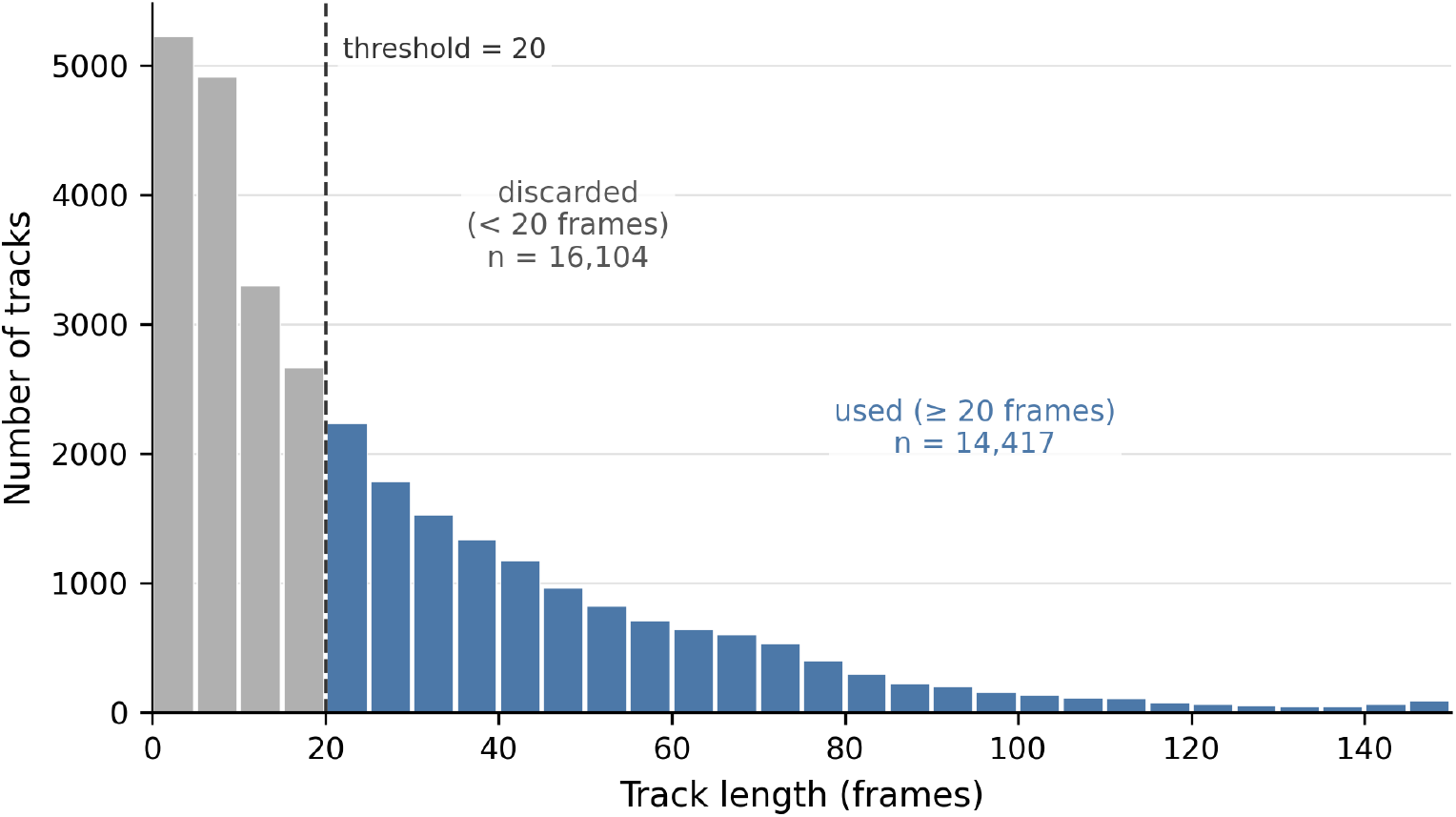
Distribution of the lengths of all 30,521 tracks obtained by tracking. Gray bars (shorter than 20 frames, 16,104 tracks) were discarded as insufficiently long, and blue bars (20 frames or longer, 14,417 tracks) were used for training and evaluation. The dashed line indicates the threshold (20 frames).

In addition, as environmental information, a 150-frame binary vessel-mask image was produced for each imaging field and used as the input to the vessel structure encoder described below. We also extracted mean intensity, SNR, and contrast from the fluorescence image data as auxiliary features.

### 2.3. Model architecture

The base model of this study is an autoregressive (AR) Transformer. Using causal self-attention, it generates a trajectory by sequentially predicting and re-encoding the cell displacement at each time point following the context. The attention blocks use a relative-position bias, with 6 heads and 6 layers. The overall data flow of the model is shown in Figure 3. The trajectory context (position, velocity, and direction of the first 10 time points) is quantized and embedded, a positional encoding is added, and the sequence is encoded by the Transformer. In the model that integrates vascular environmental information (structAR), the vessel-mask image at the same time point is encoded by a convolutional neural network and fused by time-aligned cross-attention. Finally, a mixture density head outputs the displacement distribution at each time point, and the predicted displacement is appended to the sequence to predict the next time point (autoregression).

**Figure 3.**
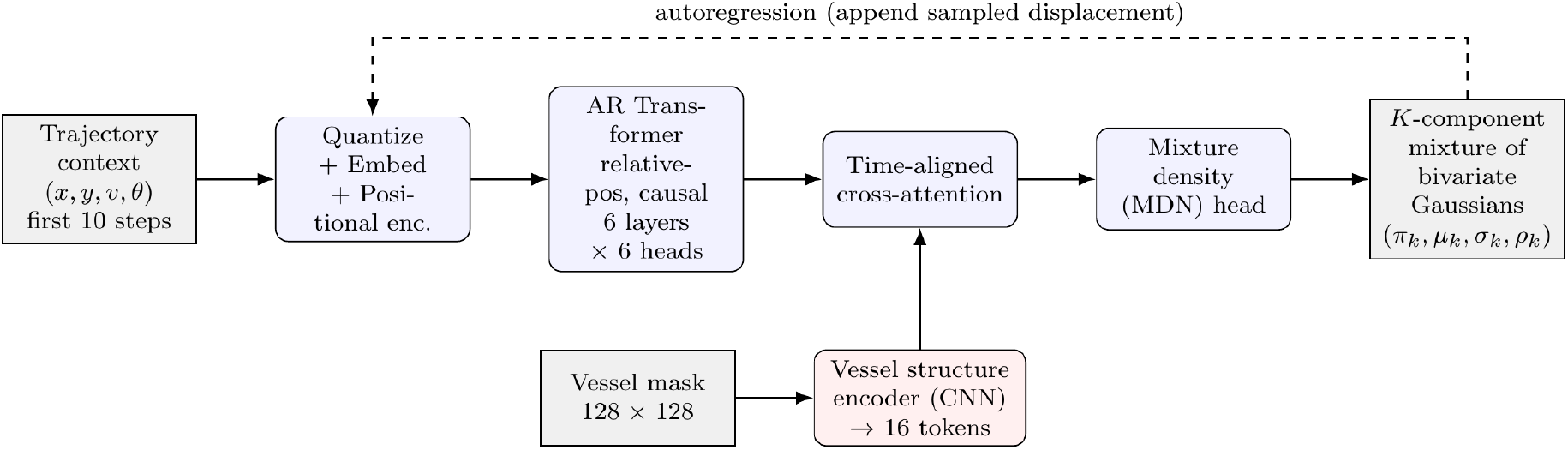
Overview of the proposed model. The trajectory context is embedded and encoded by an AR Transformer. The vessel-mask image is converted into structure tokens by a CNN (vessel structure encoder) and fused by time-aligned cross-attention (structAR). The mixture density head outputs the displacement at each time point as a *K*-component mixture of bivariate Gaussians. The dashed arrow denotes the autoregressive prediction loop.

### 2.4. Input representation

The input at each time point consists of four features: position (*x, y*), velocity, and direction. Each feature is quantized within a prescribed range and mapped to a learnable embedding (Table 2). Position is assigned a large quantization size (512) and embedding dimension (256) to preserve spatial resolution, while velocity and direction use smaller dimensions. The concatenation of the four embeddings gives a total of 768 dimensions, which is the input dimension of the Transformer.

**Table 2.**
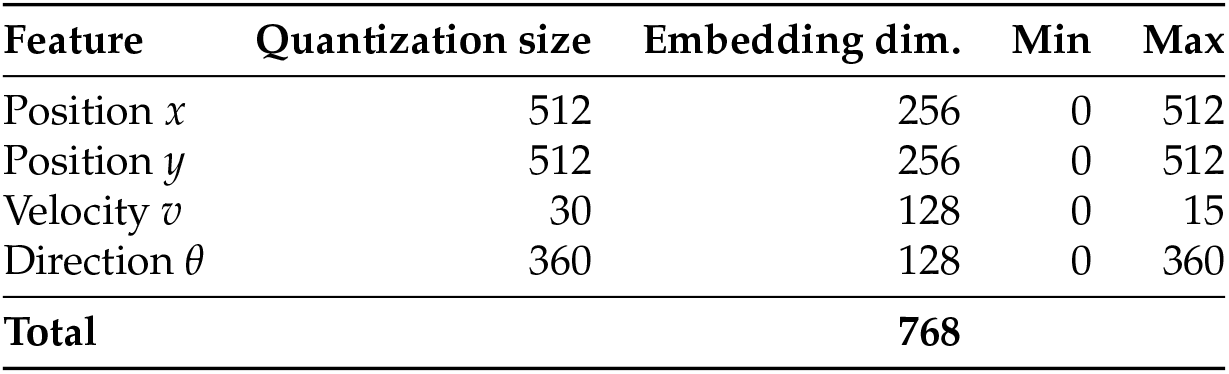
Quantization size and embedding dimension of the input features. The four embeddings are concatenated into 768 dimensions.

| Feature | Quantization size | Embedding dim. | Min | Max |
| --- | --- | --- | --- | --- |
| Position $x$ | 512 | 256 | 0 | 512 |
| Position $y$ | 512 | 256 | 0 | 512 |
| Velocity $v$ | 30 | 128 | 0 | 15 |
| Direction $\theta$ | 360 | 128 | 0 | 360 |
| **Total** |  | **768** |  |  |

### 2.5. Predictive distribution and loss (bivariate Gaussian)

The displacement (Δ*x*, Δ*y*) at each time point is output as a bivariate Gaussian distribution with mean (*µ*_*x*_, *µ*_*y*_), standard deviations (*σ*_*x*_, *σ*_*y*_), and correlation coefficient *ρ*:

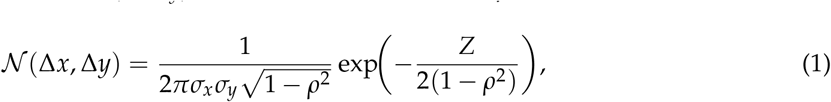

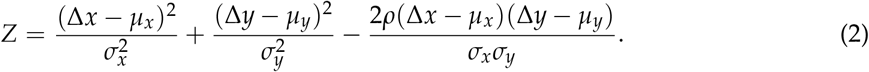

Training minimizes the negative log-likelihood (NLL) *−* ∑_*t*_ log *N* (Δ*x*_*t*_, Δ*y*_*t*_) over the valid time points. This formulation predicts not only a point estimate but also the positional uncertainty (a prediction ellipse).

### 2.6. Extension to multimodal output (mixture density)

To capture the multimodal nature of cell migration, which intermittently switches between movement (run) and rest, we extend the predictive distribution to a *K*-component mixture of bivariate Gaussians (mixture density network, MDN). The head outputs mixture weights 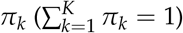 and the parameters (*µ*_*k*_, *σ*_*k*_, *ρ*_*k*_ ) of each component, so that the density is

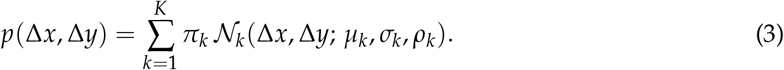

The loss is the mixture NLL, computed with a log-sum-exp for numerical stability:

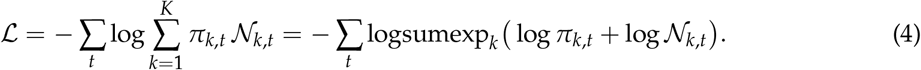

*K* = 1 is exactly equivalent to a single bivariate Gaussian. The ability to represent multimodality in a continuous displacement space without discretization is the key difference from prior grid-category approaches [7].

### 2.7. Fusion of environmental information (structAR)

In the model that integrates vascular environmental information (structAR), the binary vessel-mask image at each time point is downscaled to 128 *×* 128 and encoded by a convolutional neural network (vessel structure encoder). Five downsampling steps yield 4 *×* 4 = 16 structure tokens (64 dimensions each), reducing the cost of cross-attention. Through time-aligned cross-attention, the cell representation at each time point attends to the vessel structure tokens at the same time point, enabling predictions that inherently account for spatial context. Because the vessel mask is binary, its structure is preserved even after downscaling.

### 2.8. Training and evaluation

The data split was fixed for all experiments (random seed 42), so that every model is evaluated on the same test set. Each model configuration was trained with three random seeds (42, 43, and 44), which control the model initialization and the order of stochastic gradient descent; we report the mean *±* standard deviation over seeds and compare models through paired differences within each seed. All variants were otherwise trained under identical conditions. Optimization used Adam with weight decay, a warmup followed by cosine learning-rate decay, gradient clipping, and early stopping (patience 5). The primary evaluation metric was the validation NLL (val NLL), which does not depend on sampling; point prediction was evaluated auxiliarily by the average displacement error (ADE) and the final displacement error (FDE). For ADE and FDE, the point prediction at each step was the mean of the predicted Gaussian (for *K ≥* 2, the mean of the component with the largest mixture weight), which was fed back autoregressively, so that the evaluation is deterministic; sampling from the learned distribution was used only to visualize the predictive distribution (Figure 6). The main hyperparameters are summarized in Table 3.

**Table 3.** Main hyperparameters. The number of mixture components *K* was compared between unimodal (*K* = 1) and multimodal (*K* = 2).

| Item | Value |
| --- | --- |
| Transformer layers / heads | 6 / 6 |
| Embedding dimension | 768 |
| Vessel-mask resolution / structure tokens | $128 \times 128$ / 16 |
| Structure-token dimension | 64 |
| Number of mixture components $K$ | 1 (unimodal) or 2 (multimodal) |
| Batch size | 16 |
| Learning rate (warmup + cosine decay) | $5 \times 10^{-5}$ |
| Optimizer | Adam (weight decay 0.2, grad. clip 0.5) |
| Early-stopping patience | 5 |
| Random seeds | 42, 43, 44 (data split fixed) |
| Parameters (structAR + MDN) | $\approx 50.8$ million |

## 3. Results

### 3.1. The structure and limit of predictability

We first characterized what limits point prediction, by decomposing predictability into the movement *direction* and the activity *mode* (run/rest).

#### 3.1.1. Direction of moving cells only is predictable

The single-step movement direction was hard to predict: the alignment of velocity between consecutive steps was low (cosine similarity *≈* 0.17). However, this does not mean the direction is uniformly random. In our observation, cells intermittently switched between a diffusive resting state and a ballistic moving state (run-and-rest). When restricted to the moving population, the direction showed a consistent drift toward the brain surface (directional concentration *R ≈* 0.34, straightness 0.46; pure diffusion would give *≈* 0.16), and the direction of moving cells was predictable from the context (median angular error of the context-to-future mean velocity direction 51^*°*^, with 46% within 45^*°*^ versus 25% by chance). Crucially, a longer context did not improve the direction prediction, indicating that the predictable component of the direction has been exhausted in early steps and the remainder was irreducible. When all steps were pooled, this structure was diluted by resting cells and single-step noise, which can be the reason why the pooled direction appears uniform.

### 3.1.2. The timing of mode transitions was the bottleneck

Classifying run/rest from the recent speed yielded a sticky two-state Markov process (same-mode retention 88.6% per frame; mean dwell time *≈* 7 frames). The mode was predictable 10–20 frames ahead (autocorrelation at lag 10 is 0.22), but whether a run would persist was only weakly readable from the early context (early-to-late persistence correlation 0.10). Thus the true bottleneck of prediction was estimated not *which direction* a cell would take, but *when it started and stopping its movement*, i.e., the timing of mode transitions.

### 3.1.3. The error curve followed a persistent-random-walk law

The measured error curve of the trajectory-only model was described with high accuracy by the mean-squared-displacement (MSD) law of a persistent random walk (Fürth form; fit correlation *r* = 0.999; Figure 4). This result was a *description* by fitting, not a first-principles prediction. Attempts to reproduce the error curve from the motion statistics without fitting did not work, because the sequential memory was short (the measured velocity-autocorrelation time *τ* = 0.69 step is far shorter than the prediction horizon) and the drift did not accumulate coherently.

**Figure 4.**
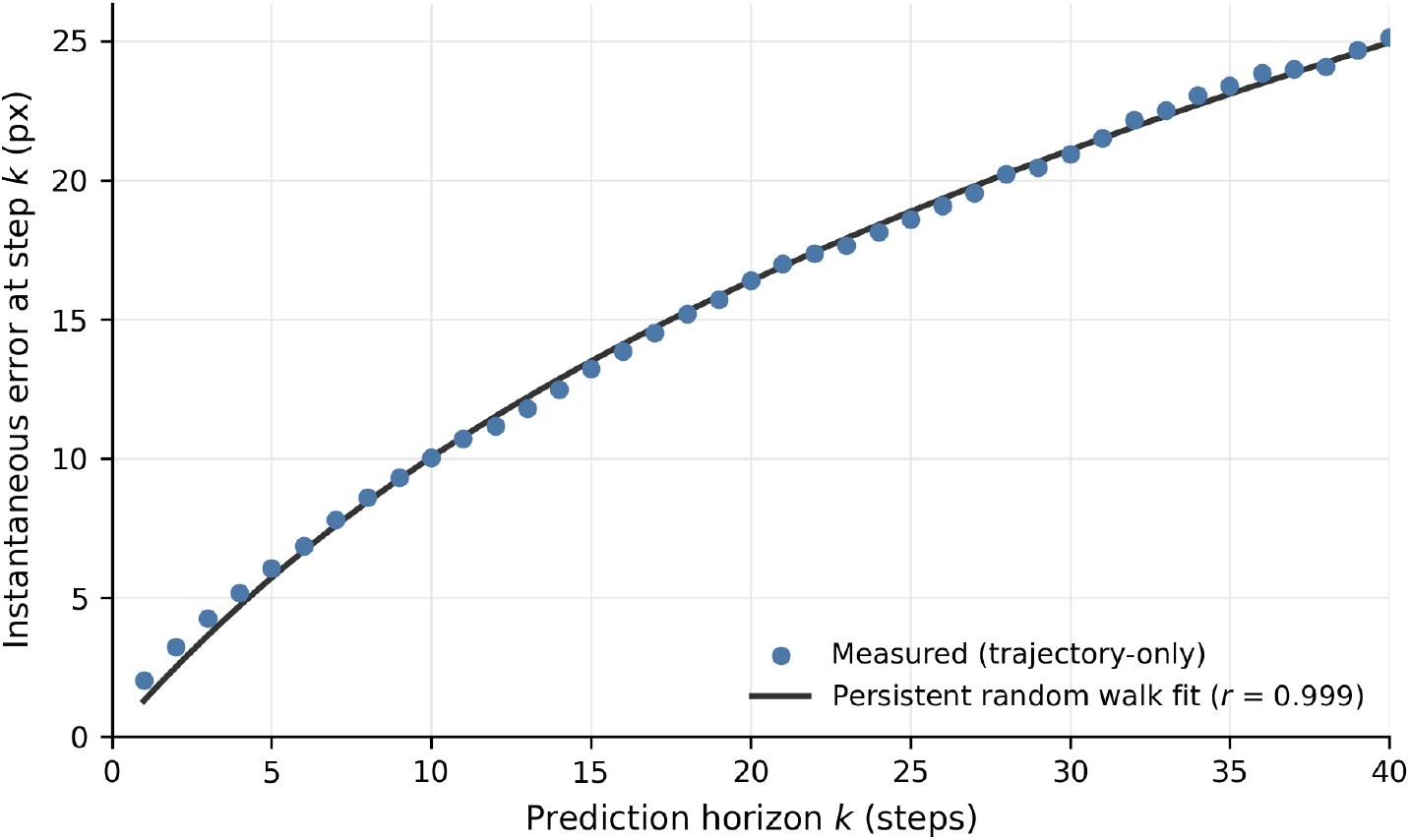
The instantaneous prediction error of the trajectory-only model as a function of the prediction horizon *k* (blue dots), together with a persistent-random-walk (Fürth MSD) fit (black line). The measured error curve is described by the fit with high accuracy (correlation *r* = 0.999), indicating that the accumulation of point-prediction error is governed by the stochastic, persistent-random-walk nature of the migration.

#### 3.1.4. Information-theoretic limit

Following the entropy-rate and Fano-inequality framework used for human mobility [9], the per-cell upper bound of predictability indicated a median of *≈* 0.47, which is far below the value for human mobility (0.93). The residuals of the best model were nearly white (lag-1 autocorrelation *≈ −*0.04; intraclass correlation 5.5%); this is consistent with the model having exhausted the predictable sequential component of the direction.

### 3.2. Multimodal output improved the predictive distribution

Motivated by the run/rest structure above, we extended the predictive distribution from a single bivariate Gaussian to a *K*-component mixture (MDN) and swept the number of components *K* (mean *±* standard deviation over three training seeds, same data split; Table 4). Almost all of the improvement occurs in the transition from a single Gaussian to two components (*K* = 1 *→* 2, NLL 4.176 *→* 3.387, i.e., a *≈* 2.2*×* improvement in likelihood), and the NLL saturated at *≈* 3.36 for *K ≥* 3 (Figure 5). These results suggest that the predictive distribution is essentially bimodal, corresponding to the run/rest two-mode structure identified above. Importantly, this improvement of the distribution had been obtained while the point-prediction accuracy (ADE) was essentially preserved.

**Table 4.** Effect of the number of mixture components *K* (trajectory-only model; mean *±* standard deviation over three training seeds, same data split). The improvement is almost entirely from *K* = 1 *→* 2 and saturates for *K ≥* 3.

| $K$ | val NLL | ADE (px) | FDE (px) |
| --- | --- | --- | --- |
| 1 (single Gaussian) | $4.176 \pm 0.003$ | $14.80 \pm 0.23$ | $25.66 \pm 0.62$ |
| **2** | **$3.387 \pm 0.005$** | **$14.44 \pm 0.10$** | **$25.13 \pm 0.33$** |
| 3 | $3.357 \pm 0.001$ | $14.96 \pm 0.52$ | $26.37 \pm 1.40$ |
| 5 | $3.357 \pm 0.002$ | $14.71 \pm 0.24$ | $25.70 \pm 0.51$ |

**Figure 5.**
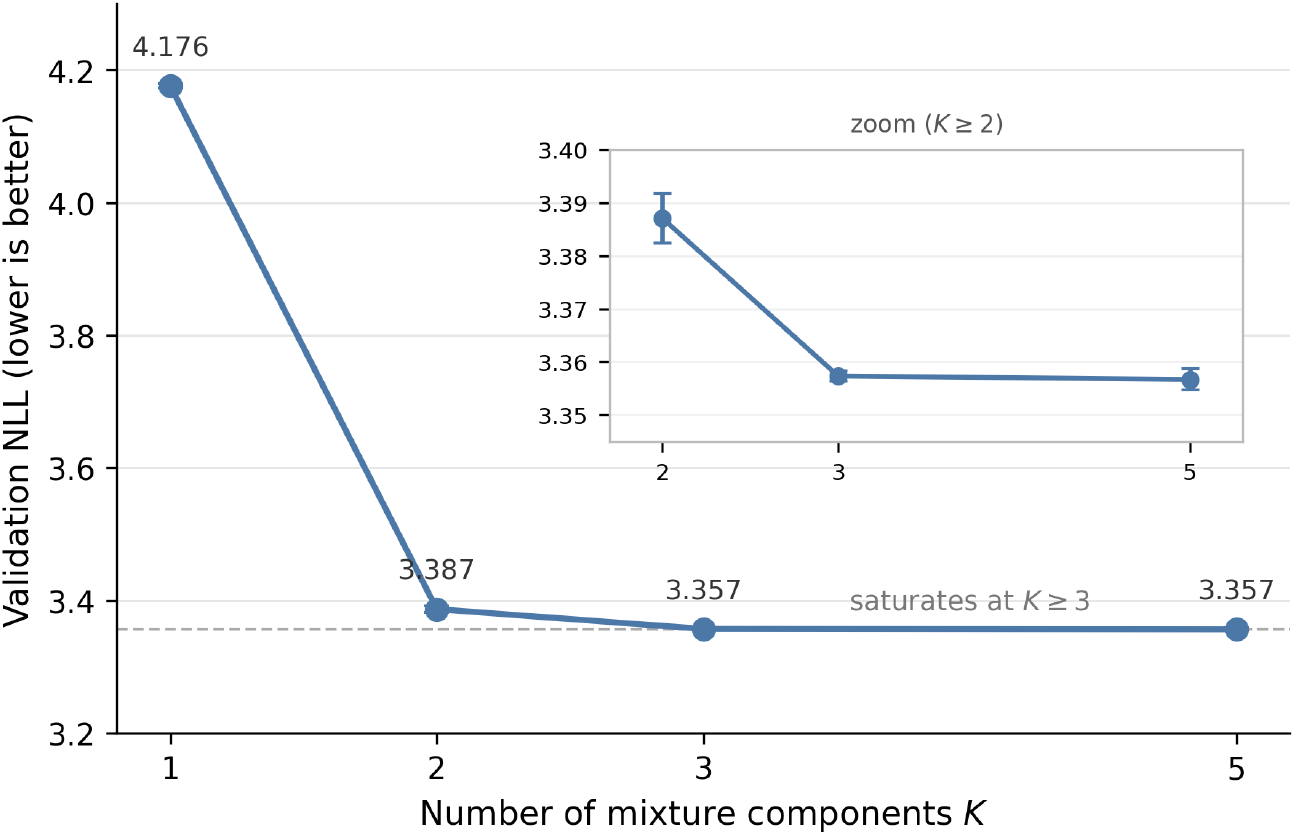
Validation NLL as a function of the number of mixture components *K* (trajectory-only model; mean *±* standard deviation over three training seeds; the error bars are smaller than the markers in the main panel). Almost all of the improvement occurs in the transition from a single Gaussian (*K* = 1) to two components (*K* = 2), and the NLL saturates for *K ≥* 3, indicating that the predictive distribution is essentially bimodal (corresponding to the run/rest structure). Inset: zoom on *K ≥* 2.

**Figure 6.**
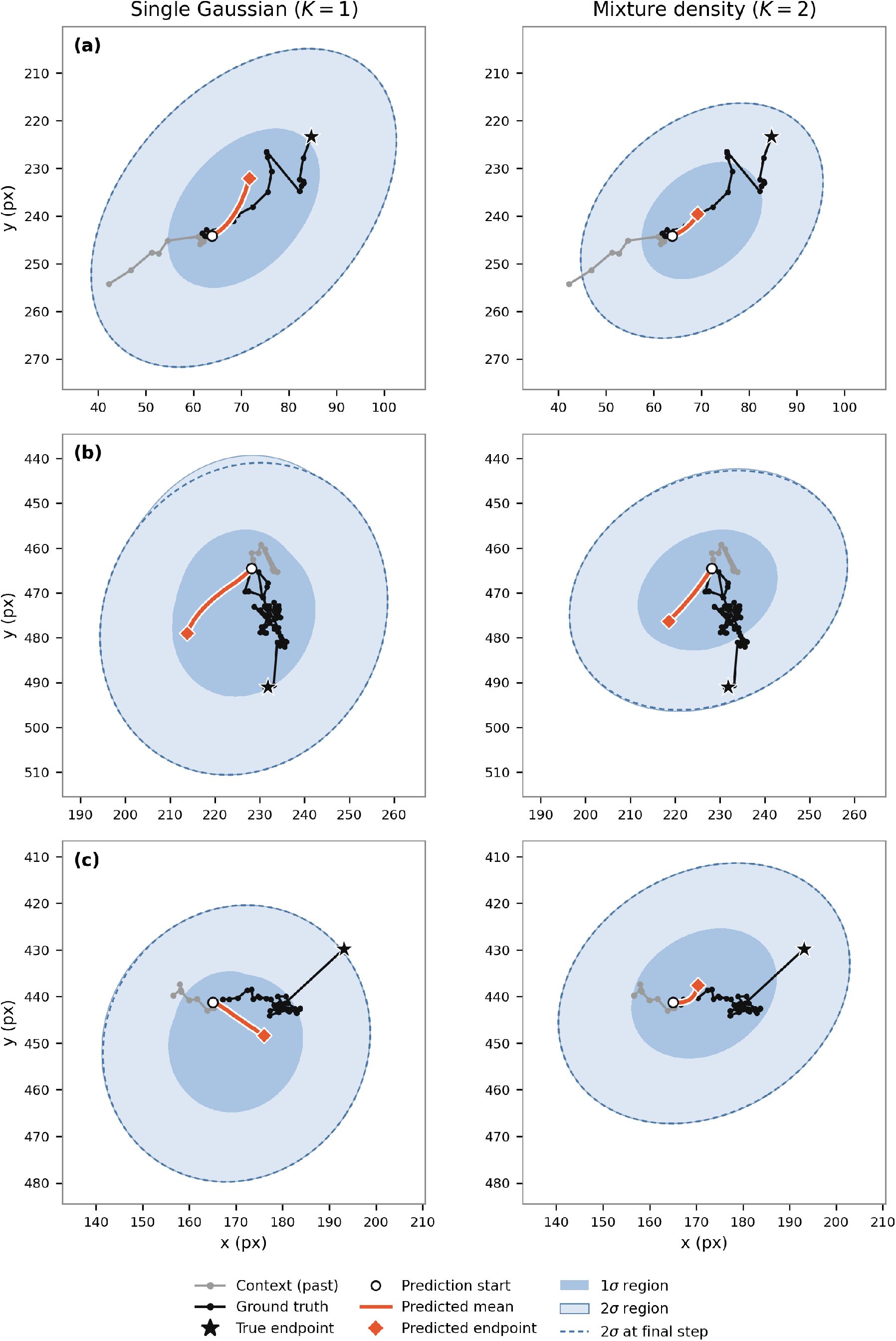
Qualitative prediction examples for three representative test cells (rows), comparing the single-Gaussian model (*K* = 1, left) and the mixture-density model (*K* = 2, right). Gray: the context (past) trajectory; black: the ground-truth future with its endpoint marked by a star; blue: the predicted uncertainty (1*σ* and 2*σ* ellipses) accumulated along the prediction horizon; dashed red: the predicted mean. The ground truth lies within the predicted distribution, which the *K* = 2 model captures slightly more tightly. The short predicted mean reflects the irreducibility of the movement direction: the model hedges its point estimate while widening the distribution.

### 3.3. Integrating environmental information yielded the best model

Finally, we integrated vascular environmental information into the multimodal model (structAR + MDN) and compared it with the trajectory-only multimodal model under the same data split and three training seeds (Table 5). The structAR + MDN model achieved the best performance on all three metrics: val NLL 3.367 *±* 0.005 (versus 3.387 *±* 0.005), ADE 13.98 *±* 0.17 px (versus 14.44 *±* 0.10, *−*3.2%), and FDE 24.33 *±* 0.52 px (versus 25.13 *±* 0.33, *−*3.2%). The improvement was obtained in every seed (paired difference in val NLL *−*0.020 *±* 0.004; paired *t*-test, *p* = 0.014), indicating a consistent, if modest, contribution of the vascular information. Its ADE and FDE were also the lowest among all the models tested, including the trajectory-only models with *K* = 3 and 5 (Table 4); evaluated by point-prediction accuracy, structAR + MDN was therefore the best model in this study (in val NLL, the trajectory-only models with *K ≥* 3 were slightly lower, 3.357).

**Table 5.**
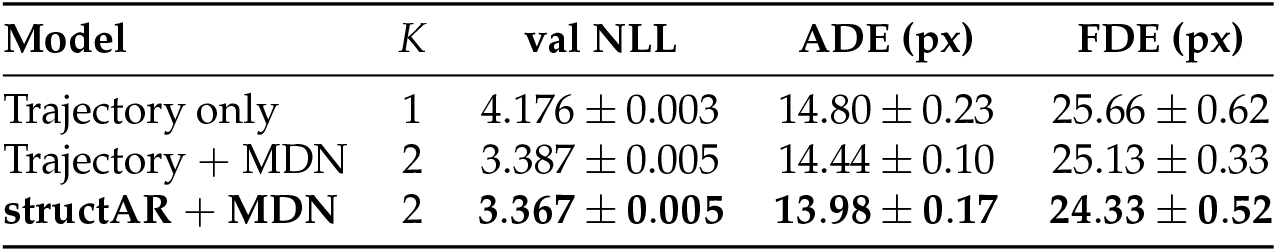
Main model comparison (mean *±* standard deviation over three training seeds, same data split). Lower is better for all metrics. The multimodal model integrating vascular information (structAR + MDN) is best on all three metrics.

| Model | K | val NLL | ADE (px) | FDE (px) |
| --- | --- | --- | --- | --- |
| Trajectory only | 1 | $4.176 \pm 0.003$ | $14.80 \pm 0.23$ | $25.66 \pm 0.62$ |
| Trajectory + MDN | 2 | $3.387 \pm 0.005$ | $14.44 \pm 0.10$ | $25.13 \pm 0.33$ |
| **structAR + MDN** | **2** | **$3.367 \pm 0.005$** | **$13.98 \pm 0.17$** | **$24.33 \pm 0.52$** |

This result is best understood together with the diagnosis above: environmental information weakly determines the activity mode (whether a cell moves) rather than the direction itself. A single Gaussian head cannot represent “which mode to take,” and therefore cannot exploit the environmental context; a mixture head, in contrast, can use the vascular context to modulate the mixture weights (mode selection). In other words, multimodal output is not only a better distribution but also the receptacle through which environmental information becomes usable for prediction.

### 3.4. Qualitative examples of the predictive distribution

Figure 6 shows representative test cells for which the predictive distribution covers the true future well. Rather than a single point forecast, the model produces a growing “forecast cone” of uncertainty ellipses (1*σ* and 2*σ*), estimated from the predictive distribution at each prediction step; the ground-truth trajectory and its endpoint (star) lied within these ellipses. Comparing the single-Gaussian (*K* = 1) and mixture (*K* = 2) models, the *K* = 2 prediction ellipses contained the true trajectory and endpoint slightly more tightly and were oriented along the direction of motion. These results are consistent with the NLL improvement (Section 3): the multimodal distribution better matches the true dispersion of the future positions. The predicted mean (dashed line) was short relative to the true displacement, reflecting the irreducibility of the direction: unable to predict the direction, the model appropriately hedged toward a small expected displacement while widening the distribution to cover the possible futures.

### 3.5. Structure of the predicted uncertainty

The predicted uncertainty *σ* was positively correlated with the cell speed (*r* = +0.512), i.e., the model expressed larger uncertainty for faster (moving) cells. On the other hand, *σ* tended to decrease along the prediction horizon (from 1.78 px at early steps to 0.96 px at late steps), suggesting that the autoregressive prediction may underestimate the uncertainty over long horizons; a quantitative calibration analysis (e.g., PIT/coverage) of the multimodal model is left for future work.

## 4. Discussion

Our results show that the predictability of astrocyte migration is not a uniform noise floor but has a definite structure that becomes visible only when the trajectory is decomposed into the movement direction and the activity mode. The single-step direction appears almost random when all steps are pooled. However, this is estimated as a dilution effect by resting cells and single-step noise, which can be the mechanism that masks the coherent, brain-surface-directed drift of the moving population (Section 3 onward). Hence, the naive conclusion that cell trajectories are unpredictable is largely an artifact of pooling heterogeneous sub-populations, reminiscent of Simpson’s paradox.

What ultimately limits prediction is therefore not which direction a cell takes, but when it starts and stops moving. The activity mode is a sticky two-state process that is predictable over 10–20 frames, yet whether a run will persist was only weakly readable from the early context. This identifies the timing of mode transitions as the true bottleneck and, at the same time, as the most promising target for future improvement.

This diagnosis also clarifies the role of environmental information. The movement patterns of the centers of cells suggest that they are more likely to undergo fine-scale movements within the vascular region, whilst the probability of large-scale movements—where the branching of blood vessels or the network geometry determines the direction of cell movement—is low. Such cellular behaviors provide less information regarding direction from the shape of the blood vessels; however, a correlation can be estimated between the distributed region and the mode of cell activities (namely, whether they are moving or not).

Consequently, a unimodal (single-Gaussian) head, which can only output a single expected displacement, cannot exploit this cue, whereas a multimodal head can use the vascular context to modulate the mixture weights, i.e., the selection between the run and rest modes. This explains why environmental information yields less improvement on its own but produces a small, consistent gain when combined with the multimodal output: multimodality is not merely a better distribution but works more as the underlying mechanism that renders environmental information usable. The diagnosis (environment informs the activity mode) and the prescription (multimodal output represents the mode) thus mesh into a single, coherent account.

Our approach is positioned within a lineage of trajectory-prediction methods. The multimodality of prediction was originally handled by TrAISFormer [7] for AIS ship trajectories through a categorical discretization of space, which had broken down at the cellular scale where displacements are only a few pixels per frame. By recovering multimodality in a continuous displacement space, our mixture-density formulation preserved point accuracy while representing the run/rest bimodality (categorical-multimodal, inaccurate to Gaussian-unimodal, accurate to mixture-Gaussian, accurate and multimodal). The contribution of this study is thus twofold: (i) a diagnosis that decomposed predictability into direction and activity mode, and located the limit in the timing of mode transitions with the support of both movement statistics and information theory; and (ii) a prescription that improved the predictive distribution without discretization and clarified the condition under which environmental information becomes useful.

Several limitations should be noted. First, the validation NLL, which does not depend on sampling, was adopted as the primary metric; across three training seeds, its standard deviation was at most 0.005, and that of ADE was 0.1–0.5 px. A preliminary comparison of models integrating multiple environmental sources (vascular and cell–cell information), trained without fixed seeds, did not show that combining sources exceeded the best single source, suggesting redundancy among the sources; this preliminary observation should be re-examined under the present protocol (fixed data split, multiple seeds, and multimodal output).

Second, the improvement obtained by integrating vascular information into the multimodal model was reproducible across three seeds but small (0.6% in NLL and 3% in ADE); confirming it with more seeds and improving the image representation (e.g., using the raw vessel intensity rather than a binary mask) is left for future work.

Third, the error curve is described by, but not predicted from first principles by, the persistent-random-walk law.

Fourth, a quantitative calibration analysis (PIT/coverage) of the multimodal predictive distribution remains to be performed.

Finally, the most important next step is to quantify the predictability of the mode transitions themselves; how far in advance the onset and termination of runs can be read, and what governs them (environment, neighboring cells, or internal state) etc., since this is the bottleneck identified here; testing whether the same direction/mode decomposition and bi- or multimodality hold for other cell types would establish the generality of the framework.

## 5. Conclusions

In this study, we predicted astrocyte migration trajectories with an autoregressive Transformer and addressed both the limit of predictability and the design of the predictive distribution. We showed that point-prediction accuracy is bounded by the irreducibility of the movement direction; we characterized this limit by decomposing predictability into the direction and the activity mode— locating the bottleneck in the timing of mode transitions—and supported it with movement statistics (a persistent-random-walk description of the error curve) and information theory (a predictability far below that of human mobility). We then showed that extending the predictive distribution to a continuous mixture of Gaussians recovers the run/rest bimodality without discretization and substantially improves the quality of the predictive distribution while preserving point accuracy, and that vascular environmental information contributes to prediction only in combination with this multimodal output. The diagnosis and the prescription are consistent through a single two-mode (run/rest) structure. Rather than chasing point accuracy beyond its intrinsic limit, this work contributes through the quality and honesty of the predictive distribution, offering a template for modeling stochastic biological processes near their limit of predictability.

## Author Contributions

Conceptualization, K.T. and N.F.H.; methodology, K.N., H.T., and K.T.; software, K.T., and K.Z. ; formal analysis, K.T.; investigation, H.T. and K.N.; resources, H.T., K.N. Y.S. and S.E. ; data curation, K.T., K.Z. and N.F.H. ; writing—original draft preparation, K.T.; writing—review and editing, N.F.H., H.T. and K.N.; visualization, K.T.; supervision, N.F.H.; project administration, N.F.H.; funding acquisition, N.F.H., H.T. and K.N. All authors have read and agreed to the published version of the manuscript.

## Funding

This research was funded by JSPS KAKENHI (JP20H05688, JP25K02437), Takeda Science Foundation, The Uehara Memorial Foundation, and Keio Gijuku Fukuzawa Memorial Fund for the Advancement of Education and Research, Keio Gijuku Academic Development Funds, and Research Promotion Organisation Fund for the Advanced AI Research Center by Kanagawa Institute of Technology.

## Institutional Review Board Statement

The original data acquisition using animals was approved by the Institutional Review Board (or Ethics Committee) of the Animal Care and Use Committee of Institute for Developmental Research, Aichi Developmental Disability Center (protocol code: 2019-13, approval date: 22 March 2019, revised on 22 November 2024).

## Informed Consent Statement

Not applicable.

## Data Availability Statement

The data presented in this study are available on request from the corresponding author. The code used for model training and analysis is available from the corresponding author on reasonable request.

## Acknowledgments

We would like to express our gratitude to Dr Donomae for his assistance in producing ground truth tracks prior to the initiation of TrackMate usage for our preliminary investigation stage. During the preparation of this manuscript, the authors used a large language model (Claude, Anthropic) to assist with drafting the text and generating figures, employed DeepL Translation and Grammarly, as well as the free version of Overleaf, for the purposes of correcting English spelling and grammar. The authors have reviewed and edited the output and take full responsibility for the content of this publication. We are deeply grateful to Dr. Penny Coggill and Prof. Viji M. Draviam for their kind and thoughtful English editing and suggestions to organise the paper.

## Conflicts of Interest

The authors declare no conflicts of interest.

## Abbreviations

The following abbreviations are used in this manuscript:

AR: Autoregressive
CNN: Convolutional Neural Network
MDN: Mixture Density Network
NLL: Negative Log-Likelihood
ADE: Average Displacement Error
FDE: Final Displacement Error
PRW: Persistent Random Walk
MSD: Mean Squared Displacement
DoG: Difference of Gaussian

## Notes

### Competing Interest Statement

The authors have declared no competing interest.

## References

1. Vaswani, A.; Shazeer, N.; Parmar, N.; Uszkoreit, J.; Jones, L.; Gomez, A.; Kaiser, L.; Polosukhi, I. Attention Is All You Need. arXiv:1706.03762v7 [cs.CL] 2023.

2. Tay, Y.; Dehghani, M.; Bahri, D.; Metzler, D. Efficient Transformers: A Survey. arXiv:2009.06732 2020.

3. Zeng, A.; Chen, M.; Zhang, L.; Xu, Q. Are Transformers Effective for Time Series Forecasting? arXiv:2205.13504 2023.

4. Lecun, Y.; Bottou, L.; Bengio, Y.; Haffner, P. Gradient-based learning applied to document recognition. Proceedings of the IEEE 1998, 86, 2278–2324. 10.1109/5.726791.

5. Krizhevsky, A.; Sutskever, I.; Hinton, G.E. ImageNet Classification with Deep Convolutional Neural Networks. In Proceedings of the Advances in Neural Information Processing Systems; Pereira, F.; Burges, C.; Bottou, L.; Weinberger, K., Eds. Curran Associates, Inc., 2012, Vol. 25.

6. Bishop, C. Mixture Density Networks. https://publications.aston.ac.uk/id/eprint/373/1/NCRG_9_4_0_04.pdf 1994.

7. Nguen, D.; Fablet, R. TrAISformer, A generatice transformer for AIS trajectory prediction. arXiv:2109.03958 [cs.AI] 2021.

8. Takahashi, K.; Zama, K.; Hiroi, N. Ship trajectory prediction using AIS data with TransFormer-based AI. In Proceedings of the 2024 IEEE Conference on Artificial Intelligence (CAI). IEEE, 2024, p. 10.1109/CAI59869.2024.00230.

9. Song, C.; Qu, Z.; Blumm, N.; Barabsi, A.L. Limits of Predictability in Human Mobility. Science 2010, 327, 1018–1021, [https://www.science.org/doi/pdf/10.1126/science.1177170]. z10.1126/science.1177170.

10. Matsumoto, K.; Azami, T.; Otsu, A.; Takase, H.; Ishitobi, H.; Tanaka, J.; Miwa, Y.; Takahashi, S.; Ema, M. Study of normal and pathological blood vessel morphogenesis in Flt1-tdsRed BAC Tg mice. Genesis 2012, 50, 561–571. 10.1002/dvg.22031.

11. Tabata, H.; Nakajima, K. Efficient in utero gene transfer system to the developing mouse brains using electroporation: Visualization of neuronal migration in the developing cortex. Neuroscience 2001, 103, 865–872.

12. Tabata, H.; Nakajima, K. Labeling embryonic mouse central nervous system cells by in utero electroporation. Develop. Growth Differ. 2008, 50, 507–511. 10.1111/j.1440-169X.2008.01043.x.

13. Tabata, H.; Nagata, K.; Nakajima, K. Time-Lapse Imaging of Migrating Neurons and Glial Progenitors in Embryonic Mouse Brain Slices. J Vis Exp 2024, 205. 10.3791/66631.

14. Tabata, H.; Sasaki, M.; Agetsuma, M.; Sano, H.; Hirota, Y.; Miyajima, M.; Hayashi, K.; Honda, T.; Nishikawa, M.; Inaguma, Y.; et al. Erratic and blood vessel-guided migration of astrocyte progenitors in the cerebral cortex. Nat Commun 2022, 13, 6571. 10.1038/s41467-022-34184-x.

15. Yusa, K.; Zhou, L.; Li, M.; Bradley, A.; Craig, N. A hyperactive piggyBac transposase for mammalian applications. Proc. Natl. Acad. Sci. USA. 2012, 108, 1531–1536. 10.1073/pnas.1008322108.

16. Zhuo, L.; Theis, M.; Alvarez-Maya, I.; Brenner, M.; Willecke, K.; Messing, A. hGFAP-cre transgenic mice for manipulation of glial and neuronal function in vivo. Genesis 2011, 31, 85–94. 10.1002/gene.10008.

17. Wang, X.; Qiu, R.; Tsark, W.; Lu, Q. Rapid promoter analysis in developing mouse brain and genetic labeling of young neurons by doublecortin-DsRed-express. J Neurosci Res 2007, 85, 3567–3573. 10.1002/jnr.21440.

18. Tabata, H.; Nakajima, K. Multipolar migration: the third mode of radial neuronal migration in the developing cerebral cortex. J. Neurosci 2003, 23, 9996–10001.

19. Kitazawa, A.; Kubo, K.; Hayashi, K.; Matsunaga, Y.; Ishii, K.; Nakajima, K. Hippocampal pyramidal neurons switch from a multipolar migration mode to a novel “climbing” migration mode during development. J. Neurosci 2014, 34, 1115–1126.

20. A., S.C.; S., R.W.; W., E.K. NIH Image to ImageJ: 25 years of image analysis. Nature Methods 2012, 9, 671–675. 10.1038/nmeth.2089.

21. Schindelin, J.; Arganda-Carreras, I.; Frise, E.; Kaynig, V.; Longair, M.; Pietzsch, T.; Preibisch, S.; Rueden, C.; Saalfeld, S.; Schmid, B.; et al. Fiji: an open-source platform for biological-image analysis. Nature methods 2019, 9, 676–682.

22. Lowe, D. Distinctive Image Features from Scale-Invariant Keypoints. International Journal of Computer Vision 2004, 60, 91–110. 10.1023/B:VISI.0000029664.99615.94.

23. Pietzsch, T.; Preibisch, S.; Tomančák, P.; Saalfeld, S. ImgLib2—generic image processing in Java. Bioinformatics 2012, 28, 3009–3011. 10.1093/bioinformatics/bts543.

